# Measuring Orthopedic Plate Strain to Track Bone Healing Using a Fluidic Sensor Read via Plain Radiography

**DOI:** 10.1101/2020.08.27.268169

**Authors:** Apeksha C. Rajamanthrilage, Md. Arifuzzaman, Paul W. Millhouse, Thomas B. Pace, Caleb J. Behrend, John D. DesJardins, Jeffrey N. Anker

## Abstract

We describe a fluidic X-ray visualized strain indicator under applied load (X-VISUAL) to quantify orthopedic plate strain and inform rehabilitative care. This sensor uses a liquid-level gauge with hydro-mechanical amplification and is visualized in plain radiographs which are routinely acquired during patient recovery to find pathologies but are usually insufficient to quantify fracture stiffness. The sensor has two components: a stainless-steel lever which attaches to the plate, and an acrylic fluidic component which sits between the plate and lever. The fluidic component has a reservoir filled with radio-dense solution and an adjoining capillary wherein the fluid level is measured. When the plate bends under load, the lever squeezes the reservoir, which pushes the fluid along the channel. A tibial osteotomy model (5 mm gap) was used to simulate an unstable fracture, and allograft repair used to simulate a stiffer healed fracture. A cadaveric tibia and a mechanically equivalent composite tibia mimic were cyclically loaded five times (0 – 400 N axial force) while fluid displacement was measured from radiographs. The sensor displayed reversible and repeatable behavior with a slope of 0.096 mm/kg and fluid level noise of 50 to 80 micrometers (equivalent to 5-10 N). The allograft-repaired composite fracture was 13 times stiffer than the unstable fracture. An analysis of prior external fracture fixation studies and fatigue curves for internal plates indicates that the threshold for safe weight bearing should be 1/5^th^ −1/10^th^ of the initial bending for an unstable fracture. The precision of our device (<2% body weight) should thus be sufficient to track fracture healing from unstable through safe weight bearing.

## I. Introduction

Following fracture fixation surgery, patients commonly ask, “Am I healing normally, and when will it be safe to resume weight bearing?” These are important questions because premature weight bearing can lead to refracture and hardware failure, while unnecessary delayed return to activities can affect quality of life. About five percent of the annual estimated 2 million fracture fixation surgeries in the United States result in delayed or impaired healing due to various reasons,[1], [2] with nearly 100,000 non-unions annually.[3]–[5] In particular, tibial fracture is the most common long bone injury, and non-union is both relatively prevalent in part due to the small tissue envelope and disabling.[6], [7] For slowly healing patients at risk of nonunion, physicians can prescribe restrictive/assisted weight bearing,[8] altered physical therapy regimes, tests to detect and address underlying metabolic disorders, apply biologics such as teriparatide,[9] prescribe electrical, ultrasound, or shockwave bone stimulation therapies, or perform revisions before hardware failure.[10]–[12] However, these are unnecessary or contraindicated in most patients, who heal normally. Unfortunately, healing timeframes depend strongly upon patient-specific factors including anatomy, age, sex, metabolic factors and comorbidities (e.g., diabetes), physical regime, and smoking.[3] Additional considerations include the fracture pattern (gap size, number of fragments, and periosteal involvement), method of treatment, hardware loosening, or implant associated infections.[13] Since no single time frame applies to all patients, individual assessments are necessary.

Currently, fracture healing is clinically monitored and managed via imaging studies, serologic markers, clinical examination, and traditional timeframes.[2] These techniques are subjective and only indirectly estimate the stage of healing.[14] The lack of quantitative measurements leads to suboptimal care and inhibits clear communication in the care team (physicians, patients, physical therapists, and physicians treating comorbidities). The need for objective measurements is widely recognized,[1], [3] for example, in a review on non-unions Hak et al. state, “The answer to whether we need a better assessment of fracture healing is an unqualified yes.”[3]

In orthopedics, fracture healing is generally considered to be the restoration of biomechanical function, especially for measures of alignment, strength and stiffness. Many studies have suggested that biomechanical measurements could improve clinical outcomes if they were easy to collect in the clinic. As a fracture heals, the fracture callus stiffens and the load to failure increases, with the two properties increasing proportionally to each other at early stages.[15] Measuring stiffness (e.g. deflection or strain for a given load) is suitable for tracking healing and risk of failure because it can be measured nondestructively (unlike load-to-failure). For tibial fractures repaired using external fixation (pins passing through the skin connected to an external plate), the fracture stiffness can be measured by applying force across the bone and measuring the resulting pin deflection or plate bending. Several clinical studies on external fixation devices found that compared to standard clinical assessment, decisions based on mechanical stiffness dramatically decreased refracture rates while also reducing average time to hardware removal.[15], [16] Similarly, in a clinical study of 27 patients with a titanium femoral plate instrumented with a strain gauge and wireless telemetry, Seide et al found a wide range of healing rates (as measured by decrease in relative elasticity), and strong correlations between mechanical and CT analysis of callus healing.[17] However, such sensors contain complex and sophisticated circuitry for sensing, power, and telemetry and have proven difficult to introduce into the market. Researchers have also proposed methods to non-invasively measure strain on orthopedic devices using ultrasound,[18] implanted magnetoelastic wireless electronic devices,[19], [20] and analysis of vibrations through the bone.[21], [22] However, all of these approaches are at early stages and would require additional infrastructure and cost to be clinically translated.

This study introduces an implantable hydraulic sensor which can easily attach to the plate to quantify tibial plate bending during the fracture healing process via plain radiography (X-ray projection imaging). Plain radiography is ubiquitously available in hospitals and routinely used to observe the hardware and fracture callus, however, it conventionally has limited value for quantification of fracture healing.[20], [23] Plain radiography can help physicians assess fracture healing and detect pathologies because of its excellent contrast for hardware/bone, low cost, extensive availability, rapidity, and lower radiation exposure compared to computed tomography (CT). However, image interpretation is qualitative, subjective, and correlates poorly with mechanical properties such as stiffness or stability.[15], [21] Previously, we made a plate attachment with a tungsten pin that moved against a scale to quantitatively indicate plate bending via plain radiography.[24] However, the 12 cm long pin was surgically cumbersome, and reducing pin length would require a protruding scale and much smaller displacement. To address this concern, herein we developed a smaller device which uses hydraulic gain to amplify the signal from plate bending and increase precision for tracking healing bone.

## II. Methods

### A. Device Operating principle

As a fracture heals and the callus stiffens, it increasingly shares load with the plate, which therefore bends less under the same load. Orthopedic plate deflection under load thus provides an objective metric for monitoring the fracture healing. Our X-VISUAL device reads plate deflection as a change in the fluid level within a channel. The 0.57 ml of fluid in the sensor is a radio-dense cesium acetate solution (85 wt % in water) and is apparent on radiographic images. Cesium acetate is a colorless, hygroscopic, ionic compound with low toxicity which has been used for medical applications and previous reports indicate that it is relatively safe for in vivo use.[22] The hydraulic action of the sensor provides mechanical gain (fluid level change/plate deflection between lever attachment and bulb) based on the ratio of the bulb’s cross-sectional area to the channel’s cross-sectional area. Fig. 1 illustrates how the sensor works. With no load applied to the fractured bone, the fluid level is fixed by the position of a lever impinging on the bulb (Fig. 1 (a)). When the bone is compressed under axial load, the fracture closes and the plate bends because the plate is off the central axis (Fig. 1 (b)). This bending action releases the stainless-steel lever away from the fluid reservoir pulling fluid from the channel and thus reducing the fluid level. Fig. 1 (b) and Fig. 1 (d) are photos of hydraulic sensor with and without load. While axial compression is applied here, the same principle would apply to axial tension or direct bending moments.

**Fig. 1.**
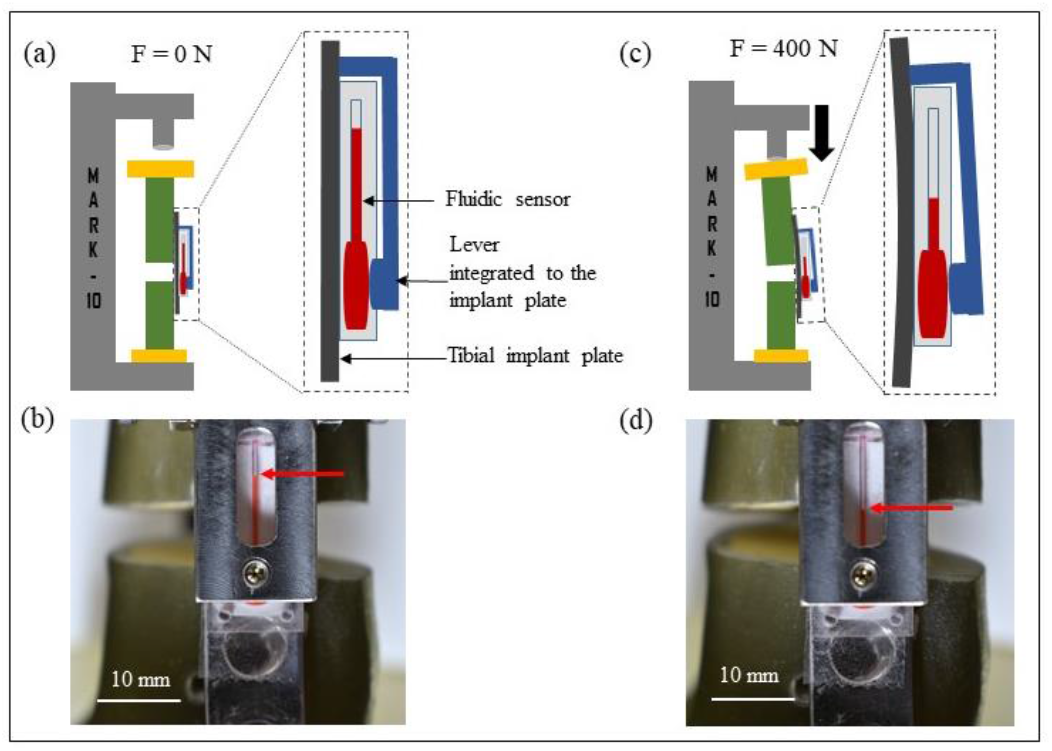
Sensor mechanism. **a)** Schematic of fluidic sensor at 0 N. **b)** Corresponding photograph in a Sawbones tibial mimic (arrow shows fluid level). **c)** Schematic of fluidic sensor at 400 N: plate bending displaces the lever, releasing the bulb and lowering the fluid level. **d)** Corresponding photograph with fluid level at 400 N.

### Device Fabrication

The passive fluidic sensor is comprised of two components, an acrylic component containing a disc-shaped fluid reservoir (bulb) attached to a millimeter diameter channel for reading fluid level, and a stainless steel mechanical lever which presses on the bulb and alters the fluid level in the channel according to degree of plate bending. Technical drawings are provided in electronic supplemental material (Fig. S1[online]). The stainless-steel lever component was designed to mount to an orthopedic plate and included an adjustable set screw to press on the fluid reservoir of the sensor. The acrylic component was machined in two halves and glued together using acrylic glue. Then, the indicator fluid was introduced into the bulb using a 27-gauge needle. Finally, the end of the channel was sealed in order to prevent escape of the fluid during operation or influx of bodily fluids.

### B. Sensor monitoring on the tibial implant plate

An unstable fracture was created at the proximal end of a Sawbones^®^ tibia composite mimic (Pacific Research Laboratories, Inc., Vashon, WA, Sawbones^®^ Model #3402). An internal fixation plate (Smith and Nephew, Lawrenceville, GA, 4.5 mm tibial locking compression plate) was employed to fix the proximal fracture. Then the hydraulic sensor was attached to the tibial plate and centered over the fracture. The lever was attached to the implant plate using screws that gripped from both sides.

### C. Measuring fluid level under load in a composite *Sawbones*^®^ tibial model

The hydraulic sensor was tested on a fractured tibia Sawbones^®^ model by monitoring the fluid displacement (mm) against the applied force (N) using an ESM-303 motorized tension/compression test stand (Mark-10 Corp., Copiague, NY) to directly evaluate the mechanical properties of the fracture site. An axial compressive load of 0 through 400 N was applied to the proximal end of the fractured model. For context, 400 N is approximately one-half body weight for an 80 kg patient. For analysis in plain radiography, the fluid reservoir was filled with cesium acetate (85 wt%).

The initial fluid level was adjusted by the turning a bottom screw attached to the lever until the “no load” level was at the top of the channel. This initial fluid level was measured in radiography, and all other fluid level displacements were calculated by subtracting this initial level. Next, the axial load on the tibia was progressively increased (loading) up to 400 N and progressively decreased (unloading) to 0 N. One set of increasing load points and one set of decreasing load points were considered as one full cycle of the study. Reproducibility of the hydraulic sensor was examined by executing five continuous loading and unloading cycles. The same procedure was carried out on a bone with an allograft (segment cut out from another Sawbones^®^ tibia) pushed into the fracture gap to simulate a repaired and reduced fracture. While such repairs are rare because traumatic fractures rarely have bone segment ejection, it simulates good fracture reduction and shows how stiffness would be expected to improve as the callus stiffens during healing. The allograft-repaired fracture was cycled over five loading cycles as with the unstable fracture.

### D. Measuring fluid level under load in a fractured human cadaveric tibial specimen

The experiments were carried out according to the Clemson University Institutional Biosafety Committee (IBC) as well as following relevant guidelines and regulations. Human cadaveric specimens were obtained via the Hawkins Foundation (Greenville, SC) from Restore Life USA (Elizabethton, TN), a nonprofit donation program, and donors had consented their body to be used for medical education and research in accordance with the Uniform Anatomical Gift Act (UAGA). The fluidic sensor was filled with radiopaque fluid in order to visualize the fluid displacement via plain radiography. The sensor was latter attached on an orthopedic implant plate in human cadaver tibia model with an unstable proximal metaphyseal tibia fracture. The mechanical lever was attached over the fluidic sensor using screws on both sides of the plate. The loading and unloading cycles and the imaging procedure were carried out as in the Sawbones composite tibial specimen above. The fractured human cadaver tibia specimen was set up with the motorized tension/compression test stand for loading/unloading experiments as shown in the figure below (Fig. 2). The tibia was loaded in compression from 10 through 400 N in 100 N increments for five consecutive loading/unloading cycles and the fluid level monitoring was done using plain radiography. Radiography was performed in Godley Snell Animal Research Center, Clemson University, Clemson, SC.

**Fig. 2.**
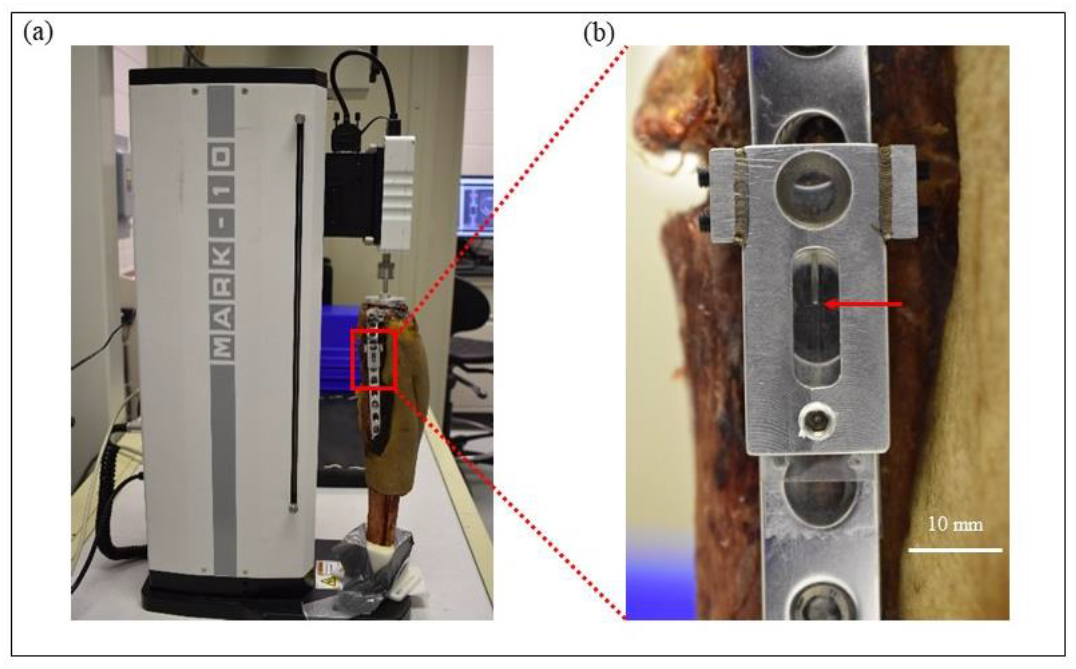
Photos of experimental setup for measuring plate strain on a cadaveric tibia with an unstable fracture. a) Fluidic sensor attached near the fracture gap. b) Zoomed-in view of the sensor (red arrow points to fluid level).

## III. Results

### A. Radiographic assessment of sensor on Fractured and allograft -repaired Sawbones^®^ tibia models under cyclic loading

A fluidic sensor filled with radio-dense cesium acetate solution (85% wt %) was used for radiographic imaging. Fig. 3 (a) shows the sequence of plain radiographs for the first of five linearly increasing and decreasing load cycles (0-400 N and back) with a Sawbones^®^ composite tibial mimic with a 5 mm osteotomy. Fig. 3 (b) shows the graphs of applied force and fluid displacement (difference from no load level) versus image number. Almost identical maximum and minimum fluid displacements were observed for each cycle indicating the reversible behavior of the fluidic sensor, with a small but consistent fluid height asymmetry at intermediate force levels indicating some hysteresis. The hysteresis curve (Fig. 3 (c)) for fractured bone has a maximum hysteresis of ~20 N and slope of 0.096 mm/kg. During loading, average displacements were ranged from 0.00 ± 0.01 mm at 0 N through 3.935 ± 0.001 mm at 400 N. The mean noise level (loading and unloading) was 0.02 mm (0.3 kg) and maximum was 0.06 mm (0.6 kg). Average displacements are tabulated and are presented in supplementary material (Table S1 [online]).

**Fig. 3.**
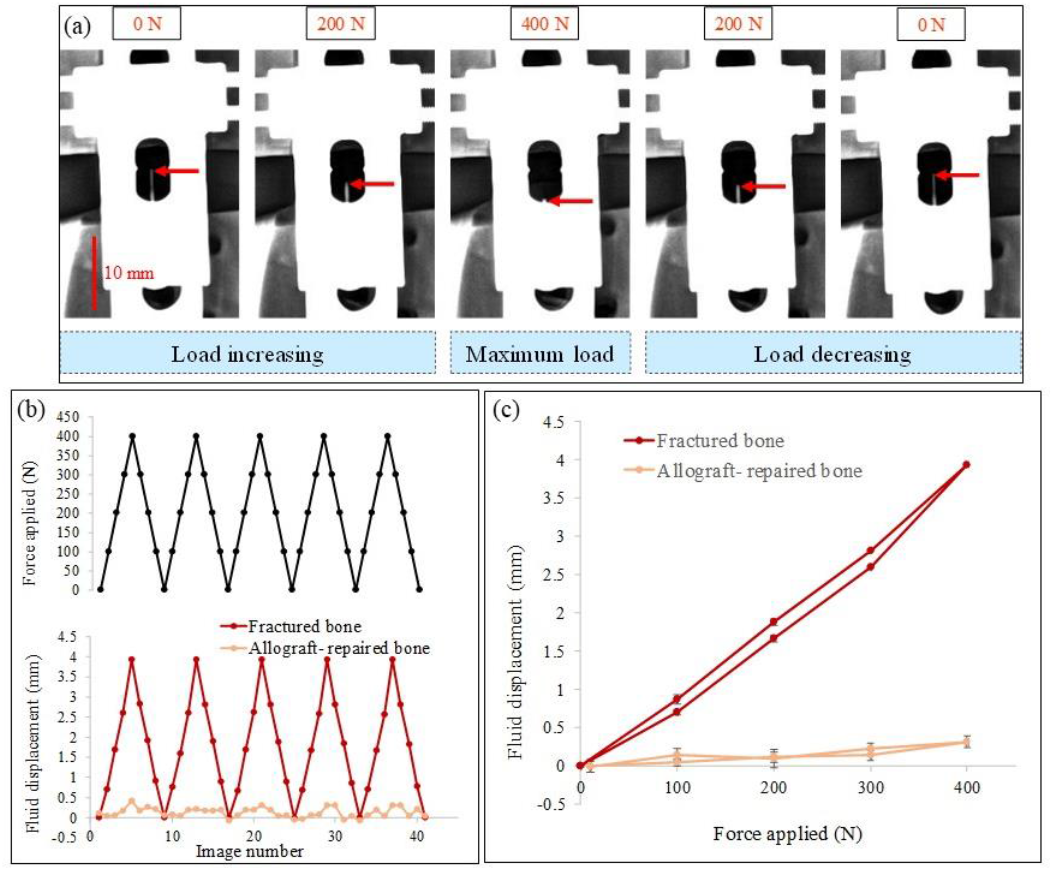
Radiographic fluid level monitoring during five loading cycles for Sawbones^®^ composite tibia model with an unstable plated fracture with/without allograft repair. a) Radiographs showing the fluid level for unstable fracture during one 0-400 N load cycle. Red arrow points to fluid level. b) Force applied and fluid displacement vs. image number for unstable fracture and allograft-repaired model. c) Average fluid displacement vs. force applied through five cycles for fractured and allograft-repaired tibia; error bars show one standard deviation from the 5 cycles.

We repeated the experiment after introducing an allograft of the same Sawbones® material to form a much stiffer construct under axial compression which simulated bone healing (or surgical interventions occasionally used to stabilize the fracture). In comparison to the fractured bone, the allograft – repaired bone showed approximately 13x less fluid displacement on radiographs with cyclic loading (Supplemental material). The fluid displacement and cyclic loading is graphed and showed in Fig. 3 (b) and 3 (c). The radiographs are shown in supplemental material (Fig. S2 [online]). The fluid displacement at 400 N (maximum load) was 0.31 ± 0.07 mm, corresponding to a slope of 0.0076 mm/kg.

### B. Sensor response on a human cadaveric tibia specimen with an unstable fracture under cyclic loading

The radiopaque fluid level changes of the fluidic sensor integrated on a fractured human cadaver tibia with an unstable fracture were monitored measured easily via plain radiography. While application of the increasing compressive loads, consistent decrease in the fluid level changes were observed. The fluid level also returned to the initial level while decreasing the compressive load. During cyclic loading, 0.9 mm of fluid displacement for 100 N was observed. Fig. 4 (a) shows the series of plain radiographs obtained at each load Fig. 4 (b) and 4 (c) are graphs showing the reproducibility of the fluid level changes for five continuous cycles. The maximum hysteresis was 0.9 mm (or ~100 N). This was larger than in the Sawbones composite due to the more complex biomechanics of the fracture, including fibula and soft tissue. During loading and unloading cycles, average displacements were ranged from 0.04 ± 0.07 mm at 0 N through 3.43 ± 0.07 mm at 400 N. The overall slope was 0.084 mm/kg (comparable to 0.96 mm/kg for the unstable Sawbones model). The mean noise level was 0.08 mm (corresponding to 1. kg), and maximum was 0.16 mm (2 kg). The average noise level was higher in the cadaveric specimen than the composite, likely due to the presence of soft tissue and more radiodense bone. However, noise was still relatively low, ~2% of displacement at 400 N.

**Fig. 4.**
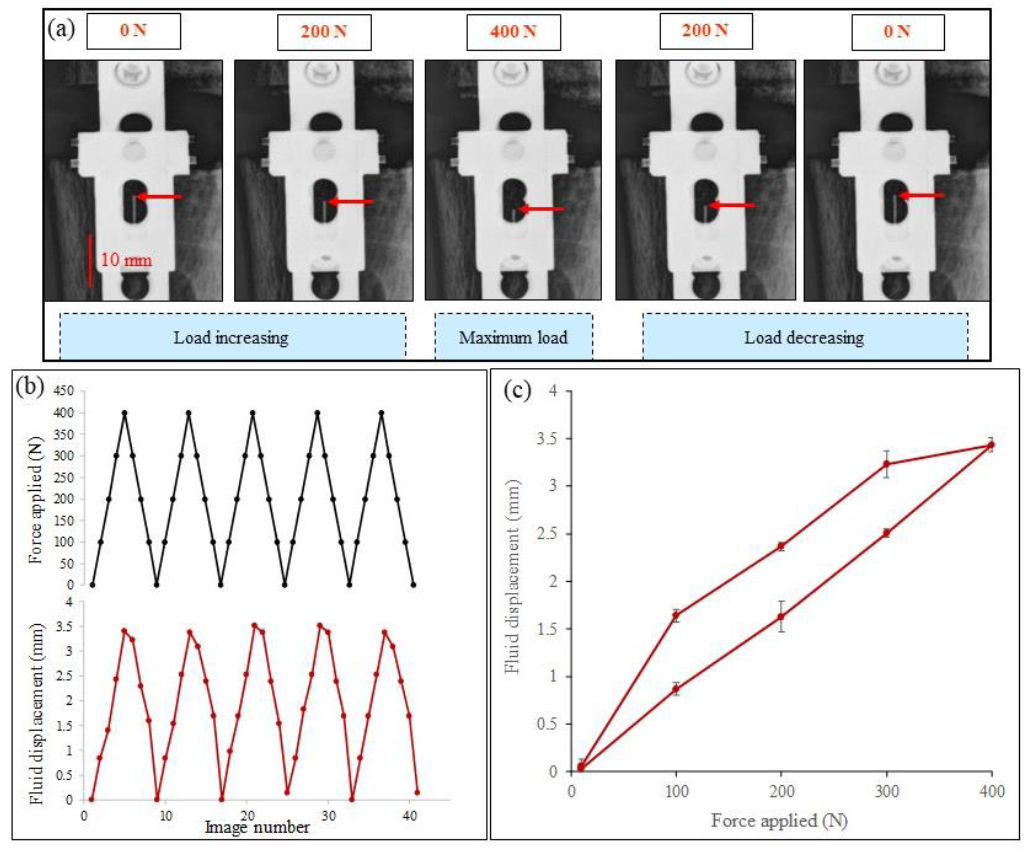
Sensor reading during five load cycles for a plated cadaveric tibia with an unstable fracture. a) Plain radiographs showing the fluid level changes with vs. applied load through one load cycle (red arrow shows fluid level). b) Force applied and fluid displacement vs. image number. c) Five-cycle-average fluid displacement vs. applied force.

## IV. Discussion

### A. Sensor Range and Precision

To be clinically useful, the sensor must be able to track strain from 0 to a maximum of bending under an unstable fracture with 1 body weight (BW), with sufficient precision to identify safe weight bearing. In the cadaver model, our sensor was able to detect displacement over the range of BW 80 μm, 2% of the displacement at 400 N (~½BW for 80 kg patient). This is over an order of magnitude better than measurements taken with standard radiography which has a typical precision of about 2-5 mm on a fracture gap, due to difficulty in identifying edges of the fracture during fracture closing, especially if there is some change in soft tissue and angle.[21], [25] Although there is no universally accepted plate strain threshold for safe weight bearing, several lines of reasoning given below suggest it should be roughly around 1/5^th^ −1/10^th^ of the initial bending during an unstable fracture, and our observed precision of 2% of the range should thus be adequate for tracking fracture healing, and readily measured the displacement for the 13x stiffer allograft-repaired model.[16], [26] More sophisticated estimates may be developed including factors relating to plate construct, healing rate, and expected activity/load, however, 2% precision will likely remain adequate. Indeed, clinically used sensors for tracking stiffness in *external* fixation devices reportedly have 3% accuracy.[27] In principle, if the range and precision are not adequate, they are adjustable in the design. The range is adjustable by the read capillary length, and the sensitivity is adjustable by controlling the length of the lever arm and ratio of chamber diameter to capillary diameter. For an unstable fracture, we observed that when the inter-fragmentary gap closed by 3 mm, the plate bending generated 4 mm of fluid displacement, which provides 1.33 of gain in inter-fragmentary motion. As the fracture heals, it would also be possible to use full BW instead of ½ which would double the fluid displacement.

#### 1) Needed range and precision based on clinical studies with external fixation devices

Clinical studies of safe hardware removal for external fixation devices (where stiffness can be directly assessed), have used several different thresholds, although they all fall in the same range. Some clinical studies have used 1/10^th^ initial bending as a threshold for device safe removal.[16] Other studies have used tibial stiffness between 8.5 – 20 Nm/°, with 15 Nm/° most common, which is about 25% of the stiffness of intact bone. Indeed, Richardson found that exclusive use of this threshold to decide about when to remove hardware allowed removal of external fixation allowed patients to remove devices 2.3 weeks earlier on average compared to traditional assessment without strain measurements, while longer retention of the fixation in slowly healing patients decreased re-fracture rates from 7% to 0%.[26] The 15 Nm/° threshold would correspond to 1/5^th^ of the bending we observe for our unstable plated fracture in Sawbones and cadaveric models. If external fixation hardware can safely be removed when it is carrying 1/5^th^ to 1/10^th^ of the load applied to the tibia, logically we expect at these thresholds (or perhaps even earlier) it would be safe to bear weight for internally fixed fractures where the hardware is retained.

#### 2) Needed range and precision based on animal and clinical studies with instrumented internal fixation devices

Several animal and human studies have measured strain during healing. In a sheep study with instrumented plates (rosettes of strain gauges) read using percutaneous wires show increase of the total plate surface strain under bending for different gait speeds (2-5 km/h) found that the strain decreased by a factor of about 7-8 as the fracture heals in the first 6 weeks, and stabilized for the last two weeks.[18] Similarly, in a clinical study of 27 human patients with a titanium femur plate instrumented with a strain gauge and wireless telemetry, Seide and co-workers found a wide range of healing rates (as measured by decrease in relative elasticity), and strong correlations between mechanical and CT analysis of callus healing.[17] Stiffness of 30% corresponded to bridging bone up to the intramedullary canal by CT, and ~10% of initial correlated to bridging throughout the femur by CT. For all these reasons, a stiffness of 1/5^th^ to 1/10^th^ of initial unstable fixation is likely to be indicative of safe weight bearing, and a noise level of 1-3% of 400 N (½ BW) as observed in our fluidic sensor should be comfortably able to measure this stiffness level.

#### 3) Needed range and precision based on plate fatigue

An analysis based on fatigue of plates gives a similar 1/10^th^ to 1/5^th^ threshold, although it suggests that thresholds could be modified based on activity level, implant fatigue, implant placement, and initial bending. Although no single study presents the entire applied load vs. average or probable cycles to failure, from prior studies at low and high cycle number we can estimate its shape for bent and unbent plates based on three studies.[28]–[30] Plates are designed to align bone fragments and limit interfragmentary motion, but allow some bending to encourage callus formation.[31] Tibial plates support short term weight bearing above 1×BW, but will fail under larger loads and eventually fatigue under repetitive cycling of 1×BW. For a single cycle, *Lindeque et al* found that proximal locking compression plates (LCP) failed under loads of 1720 N for Synthes plates, and Xu et al found Zimmer plates and 1500 N for mid-tibial shaft LCP, averaging to around 2x BW for an 80 kg person.[28], [29] Additionally, *Brunner et al* studied high cycle fatigue failure for normal and bent titanium plates (plate bending is sometimes done deliberately to better fit the bone contour and also can occur naturally during the fixation process). At 106 cycles in NaCl solution they report 7 Nm for unbent plates and 3 Nm for plates that were bent prior to fixation.[30] Assuming that the load was axial and that the fracture was treated midshaft where the tibia radius is ~18.2 mm[32], these correspond to 732N, and 323N, respectively. Fig. 5 (a) plots these values, and further assumes that the curve looks approximately s-shaped with constant failure at low cycle-number (1-100), followed by high cycle fatigue characteristics between 10^3^ and 10^6^ cycles from Bunner. The ratio of low single cycle yield strength to the endurance limit for high cycles with the unbent plate is ~1.8, and is similar to ASM value of ~1.9.[33] From Brunner’s work, the bent plate appears to have ~200 N lower strength at low cycle number, and around twice as much at high cycle number. While these are rough estimates, and more sophisticated studies can be performed for specific plates, amount of pre-bending, and position on the bone, they capture the general characteristics of fatigue.

**Fig. 5.**
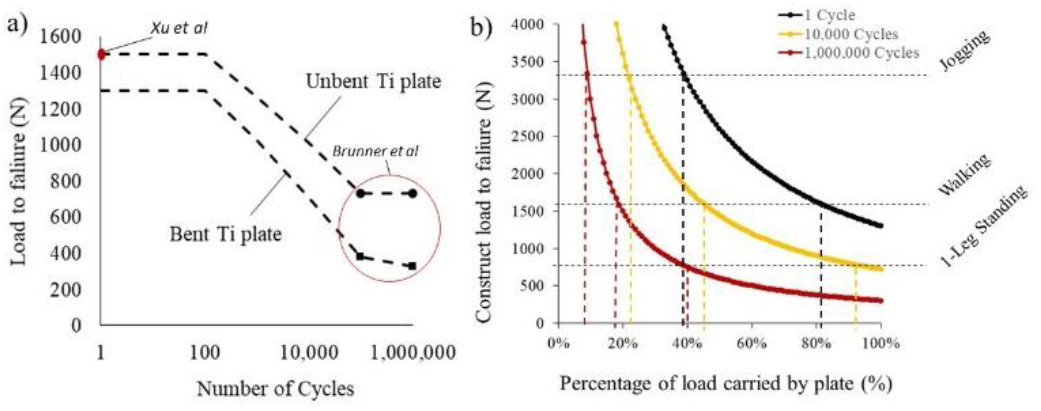
Plate load to failure for different activity levels. a) Estimated load (N) to failure vs. number of cycles for bent and unbent plates. b) Construct load to failure (N) and associated activity vs. percentage of load carried by plate assuming bent plate from part (a) and 80 kg patient.

Using the average load vs. cycles curve Fig. 5a, we can estimate how much load would be tolerated for a given number of cycles and fraction of load is carried by the implant (with rest shared by the fracture callus). Figure 5b shows this analysis for three loads: 1 cycle, 10,000 cycles (typically accepted as initial postop rehabilitation period[34]–[36], and 10^6^ cycles (close to the endurance limit). A bent plate from fig. 5a was chosen to be conservative and because plates often bend upon implantation; an analysis based on the unbent curve is shown in Supporting Information Figure S3. For single cycles (black curve), orthopedic plates can comfortably support the load from a 80 kg person standing on one leg with a completely unstable fracture (~784 N) but will probably fail during normal walking (2.5-2.8 ×BW) or jogging at 5 mph (4.2×BW).[37] As the fracture callus stiffens, it shares an increasing fraction of the load, and when it carries 50% of the load, peak loads from short term walking would be safe; however, a normally active person takes over 10^6^ steps/year, and after 10^6^ steps, the load to failure for many plates decreases to about 25% of failure load for a single cycle.[38] For safe weight bearing after 10^6^ walking steps one would need the load on the plate to be around 1/5^th^ to 1/6th of 1 BW, with the callus taking the rest, and 1/8th to 1/9th for jogging. This line of reasoning suggests that a heavier and more active patient may have a more conservative threshold (or be fitted with a stiffer plate), or that a rapidly healing patient with less fatigue may allow earlier activity.

### B. Comparison to other loaded X-ray techniques

Our sensor design improves upon existing techniques to measure fracture displacement load using X-ray imaging which are either not sufficiently precise or not readily adaptable to clinical application. Although X-ray images can show the hardware and fracture callus, they do not measure the mechanical properties of the fracture.[5], [6] With certain fracture types, X-ray images are taken with and without weight-bearing to measure the relative motion of bones in order to assess fracture stability. However, this procedure has proven insufficiently sensitive, even in cases such as spine fusion, where spinous processes are clearly evident and can move millimeters.[7]–[9] For example, Song et al. found interobserver variation in measuring spinal process motion to be 1.5 mm (95% confidence interval), an order of magnitude greater than our fluid level precision.[39] Quantitative motion analysis (QMA) software of the intervertebral motion only slightly improves upon manual radiograph analysis.[3] Lower extremity fracture gap movement is even more difficult to access (e.g., ±5 mm for acetabular fractures),[10]–[13] and the fragments move less.

Use of clear markers dramatically improves precision. For example, in radiostereometric analysis (RSA) several tantalum beads are surgically inserted near the fracture site and X-ray images are simultaneous taken at two angles to triangulate the bead position. Precision is typically within ~20-50 μm and bead motion under a load is then used to track fracture stiffness and healing.[40] However, RSA is clinically cumbersome as it needs specialized instruments, increased surgical time/cost to introduce tantalum beads, expertise to perform procedure and analyze results, and the analysis can potentially be confounded by bead placement and migration.[40]

Our hydraulic sensor precision is high compared to direct measurements of fracture gap displacement due to a combination of the hydraulic gain and measuring a clear and unambiguous sensor fluid level. In our experiments, we found average cycle-to-cycle fluid levels variation of 20-70 μm in sawbones, and 80 μm in cadaveric models. This variation could come from a combination of changes in mechanical loading conditions (e.g. settling), and from uncertainty in reading the radiographs (noise). Since the readings do not appear to systematically drift in time, noise is likely the main source of variation, and assuming the variation is noise sets an upper limit to the noise level. A precision of 80 μm is similar to the 100 μm interobserver variation we found in measuring displacement of radiopaque tungsten wires in a pH sensor based on pH-responsive hydrogels swelling[41]. It is somewhat worse than RSA analysis of tantalum beads (20-50 μm), but an order of magnitude better than direct fracture gap motion analysis with plain film X-rays.

The fluidic sensor can be easily attached to orthopedic plates during surgery, preop (or eventually integrated into plates). Our group previously developed a pin indicator attached to the side of the plate which moving against a scale to provide quantitative readings of plate bending.[24] The gain (pin motion/maximum interfragmentary motion) was approximately the ratio of the pin length to the bone diameter. However, the pin length was 12 cm which was cumbersome. In contrast, the fluidic sensor has hydro-mechanical gain (fluid level displacement/lever displacement during plate bending) almost equal to the fluid reservoir area divided by the capillary channel cross sectional area. In our case, the diameter of the reservoir is 7 mm (area 49 mm^2^) and the fluid channel diameter is 0.75 mm (area 0.56 mm^2^) providing a gain of somewhat less than 87.5 (less because the lever pushes on the edges of the reservoir less than the center). This gain allowed us to use a much shorter sensor (2.5 cm instead of 12 cm), which gives it a smaller profile, makes it easier to use, and measures bending of the specific section of the implant.

The sensor attaches to the implant using simple screws to grip the side of the plate and could be mounted prior to or after fixing the plate to the bone. In future, simpler attachments could be used, or sensors could be integrated into the plate. A screw attached to the lever was used to adjust the initial fluid level to account for any plate bending during implantation; in future, longer fluid channels along the side of the plate could be used to increase the maximum sensor range. Overall, the fluid displacements provide straightforward quantitative measurements of the plate strain, thus adding more information to the plain radiographs which are already been used in routine patient workups.

### C. Limitations

While these studies have shown the principle of a fluidic X-VISUAL sensor, much work remains before it can be clinically applied. First, the design size and shape need to be optimized. The thickness of fluidic sensor together with mechanical lever add height over the orthopedic plate which could be inconvenient for the patient. Thus, in future, the fluidic sensor and the mechanical lever may be miniaturized and placed along the side of the plate, where there is more room. Third, we estimated the threshold for safe weight bearing would likely be between 1/5^th^ and 1/10^th^ initial unstable bending, however, this would need to be verified and patient specific variables may need to be considered. In addition, the sensor was tested only on a tibial plate, and translation to other plates and devices would require site-specific optimization of the sensor size, shape, and gain. Finally, unlike devices using wireless radio-telemetry, our device is read by X-ray imaging, which would permit only periodic measurements during visits to a medical facility where loaded/unloaded radiographs are acquired.

## V. Conclusions

The fluidic sensor enables monitoring orthopedic plate bending directly and quantitatively with higher resolution compared to the common methods available today. Noise levels of 80 μm or less (corresponding ~1 kg of body weight passing through the plate) which is likely sufficient to help physicians quantify the plate bending therefore callus stiffness during fracture healing. Such information would help physicians determine when the patient is safe to start weight bearing to better communicate rehabilitation protocols with patients and their medical team.

## Acknowledgment

This work was funded by National Institute of Health (NIH) under COBRE grant 5 P20 GM103444-07. Radiography was performed in Godley Snell Animal Research Center, Clemson University, Clemson, SC. Human cadaveric specimens were obtained via the Hawkins Foundation (Greenville SC) from Restore Life USA (Elizabethton, TN), a nonprofit donation program, and donors had consented their body to be used for medical education and research in accordance with the Uniform Anatomical Gift Act (UAGA).

## Notes

### Competing Interest Statement

JNA, CJB, and JD are inventors on related US Patent US10667745B2 and founders of Aravis BioTech LLC, which has licensed the rights to the patent from Clemson.

