## Supplementary material for "Measuring Orthopedic Plate Strain to Track Bone Healing Using a Fluidic Sensor Read via Plain Radiography": Figure S1. Figure S2. Figure S3. Table S1. Table S2. Table S3.

**Contents:**

Figure S1. SolidWorks™ designs of two major parts of the sensor Page 2

Figure S2. Series of plain radiographs of the allograft-repaired bone under load Page 2

Figure S3. Plate load to failure for different plates and activity levels Page 3

Table S1. Average displacements for Sawbones tibia mimic with an unstable fracture Page 4

Table S2. Average displacements for Sawbones tibia mimic with an allograft-repaired fracture Page 4

Table S3. Average displacements for cadaver tibia with an unstable fracture Page 4

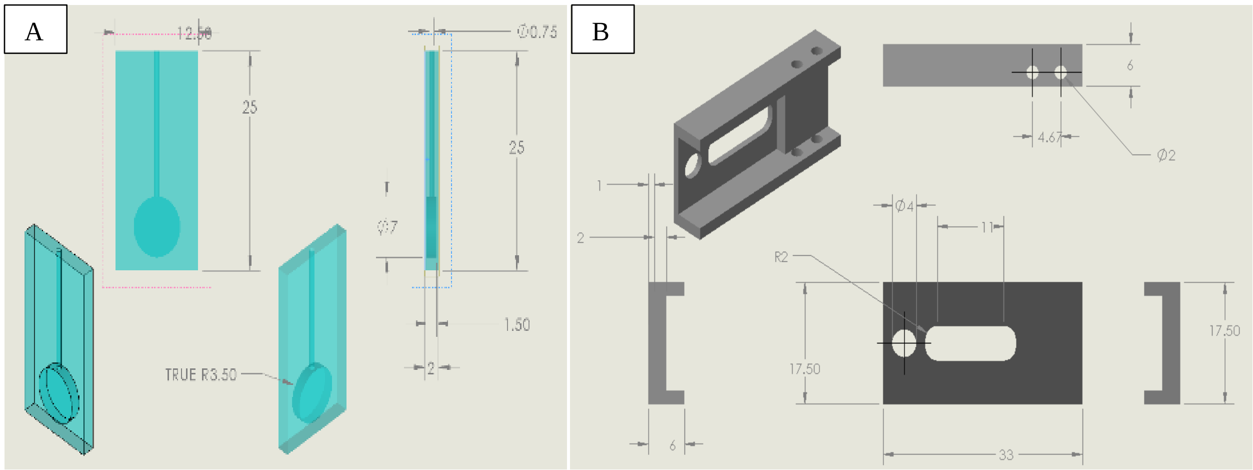


**Fig. S1.** SolidWorks™ designs of two main parts of the hydraulic sensor showing different views of the designs **A)** bulb and fluid capillary sensor; **B)** lever part of the sensor with one end which attaches to the plate, and a second end which presses on the bulb with a pressure and displacement which depend on degree of plate bending. (all measurements are in mm)

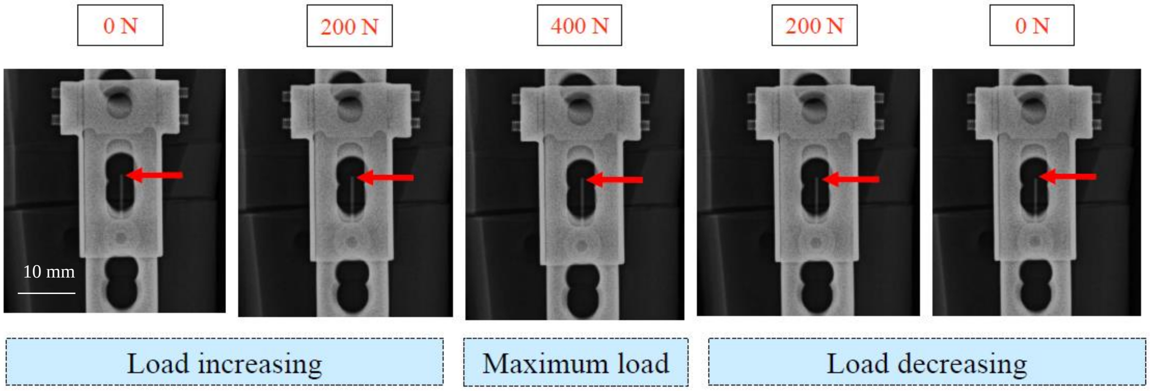


**Fig. S2.** Plain radiographs showing the fluid level changes (little or no change) for allograft- repaired tibia with respect to applied load through one cycle. (Red arrow points to fluid level)

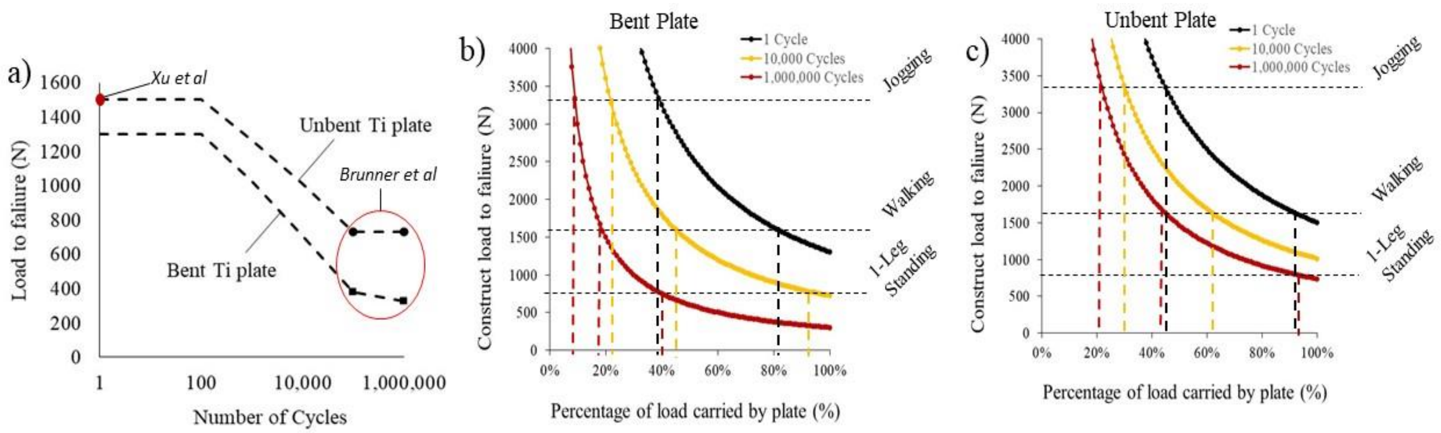


**Fig. S3.** Plate load to failure for different plates and activity levels; a) Estimated load (N) to failure vs. number of cycles for bent Ti plates and unbent plates. b) Construct load to failure (N) and associated activity vs. percentage of load carried by plate assuming bent plate from part (a) and 80 kg patient. c) Construct load to failure (N) and associated activity vs. percentage of load carried by plate assuming unbent plate from part (a)

**Table S1.** Average displacements at different loads for *Sawbones*® tibia models with an unstable fracture

| Loading | | Unloading | |
|---|---|---|---|
| Load applied (N) | Displacement (mm) | Load applied (N) | Displacement (mm) |
| 0 | 0.00 ± 0.01 | 400 | 3.935 ± 0.001 |
| 100 | 0.70 ± 0.04 | 300 | 2.82 ± 0.01 |
| 200 | 1.66 ± 0.04 | 200 | 1.88 ± 0.04 |
| 300 | 2.60 ± 0.02 | 100 | 0.87 ± 0.06 |
| 400 | 3.935 ± 0.001 | 0 | 0.00 ± 0.01 |

**Table S2.** Average displacements at different loads for *Sawbones*® tibia models with an allograft-repaired fracture

| Loading | | Unloading | |
|---|---|---|---|
| Load applied (N) | Displacement (mm) | Load applied (N) | Displacement (mm) |
| 0 | 0.00 ± 0.09 | 400 | 0.31 ± 0.08 |
| 100 | 0.04 ± 0.04 | 300 | 0.23 ± 0.07 |
| 200 | 0.11 ± 0.07 | 200 | 0.10 ± 0.12 |
| 300 | 0.14. ± 0.06 | 100 | 0.14 ± 0.08 |
| 400 | 0.31. ± 0.08 | 0 | 0.00 ± 0.07 |

**Table S3.** Average displacements at different loads for Human cadaver tibia with an unstable fracture

| Loading | | Unloading | |
|---|---|---|---|
| Load applied (N) | Displacement (mm) | Load applied (N) | Displacement (mm) |
| 0 | 0.03 ± 0.06 | 400 | 3.43 ± 0.07 |
| 100 | 0.87 ± 0.06 | 300 | 3.23 ± 0.14 |
| 200 | 1.63 ± 0.16 | 200 | 2.37 ± 0.04 |
| 300 | 2.51 ± 0.04 | 100 | 1.64 ± 0.07 |
| 400 | 3.43 ± 0.07 | 0 | 0.05 ± 0.07 |
